# Using the Wax moth larva *Galleria mellonella* infection model to detect emerging bacterial pathogens

**DOI:** 10.1101/327015

**Authors:** Rafael J. Hernandez, Elze Hesse, Andrea J. Dowling, Nicola M. Coyle, Edward J. Feil, Will H. Gaze, Michiel Vos

## Abstract

Climate change, changing farming practices, social and demographic changes and rising levels of antibiotic resistance are likely to lead to future increases in opportunistic bacterial infections that are more difficult to treat. Uncovering the prevalence and identity of pathogenic bacteria in the environment is key to assessing transmission risks. We describe the first use of the Wax moth larva *Galleria mellonella*, a well-established model for the mammalian innate immune system, to selectively enrich and characterize pathogens from coastal environments in the South West of the U.K. Whole-genome sequencing of highly virulent isolates revealed amongst others a *Proteus mirabilis* strain carrying the *Salmonella* SGI1 genomic island not reported from the U.K. before and the recently described species *Vibrio injenensis* hitherto only reported from human patients in Korea. Our novel method has the power to detect novel bacterial pathogens in the environment that potentially pose a serious risk to public health.

## Introduction

Emerging infectious diseases (EIDs) pose a major threat to human health ^1^. A large proportion of EIDs are caused by bacteria (estimated to be 54% ^1^ and 38% ^2^). Although most emerging bacterial pathogens have zoonotic origins, a large proportion of infectious bacteria are free-living, for instance being associated with food ^3^, drinking water ^4^ or recreational waters ^5^. Microbial safety is routinely assessed through the quantification of Faecal Indicator Bacteria (FIB) ^6^. However, many FIB lineages are not associated with disease and there is no *a priori* reason to expect a relationship between FIB abundance and non-gastrointestinal disease (e.g. ear or skin infections). There are dozens of bacterial genera occurring in natural environments that are not primarily associated with human or animal faecal contamination but that are able to cause opportunistic infections (e.g. ^7^). Alternatives to FIB such as quantification of pathogen-specific genes via molecular methods ^8^, flow cytometry (e.g. ^9^) or isolation of specific pathogens (e.g. ^10^) either are not linked to infection risk, are based on costly methodologies or are limited to a subset of ‘known knowns’. The current lack of a direct screening method for the presence of pathogenic bacteria in environmental samples is therefore a major barrier to understanding drivers of virulence and ultimately infection risk.

We demonstrate the use of the Wax moth larva *Galleria mellonella* as a bioindicator for microbial water quality, and a means to selectively isolate and characterize pathogens. *G. mellonella* is a well-established model system for the mammalian innate immune system and has been used extensively to test for virulence in a range of human pathogens by quantifying survival rate after injection of a defined titre of a specific strain or mutant ^11^. Bacterial virulence in *Galleria* is positively correlated with virulence in mice ^12^ as well as macrophages ^13^. Instead of quantifying the virulence of a specific bacterial clone, here we measure *Galleria* survival after injection with entire microbial communities from concentrated environmental water and sediment-wash samples. We isolate bacteria responsible for *Galleria* mortality and assess their pathogenic potential through whole-genome sequencing.

## Results

Our survival assay shows that *Galleria* mortality after 72 hours varied widely between both water and sediment samples collected at two dates from eight locations across Cornwall (U.K.), ranging from 5% to 95% (Fig. S2). Injection of buffer solution or filtered (0.22 μm) samples yielded zero mortality, demonstrating that injection was not harmful, and that samples did not contain lethal concentrations of pollutants or toxins. Mortality was largely congruent with FIB counts as well as total bacteria density (as quantified by flow cytometry and total viable counts on LB, Fig. S3), although there was substantial variation (Fig. S4 and Supplementary Results).

We chose four environmental samples exhibiting high (≥ 70%) *Galleria* mortality to isolate the clone(s) responsible for infection and reinoculated these samples to isolate bacteria from the haemocoel of infected, freshly killed larvae. All samples yielded a single colony type on each agar type, indicating that infections were (largely) clonal. A single clone was picked for each sample, grown up and assayed using three inoculation densities (1×10^2^ CFU, 1×10^4^ CFU, and 1×10^6^ CFU) (Fig. 1). All clones displayed high levels of virulence and were characterized using whole-genome sequencing (Fig. 1). We specifically focused on the identification of virulence- and antibiotic resistance genes (ARGs) as compiled in the VFDB ^14^ and CARD ^15^ databases respectively.

**Figure 1.**
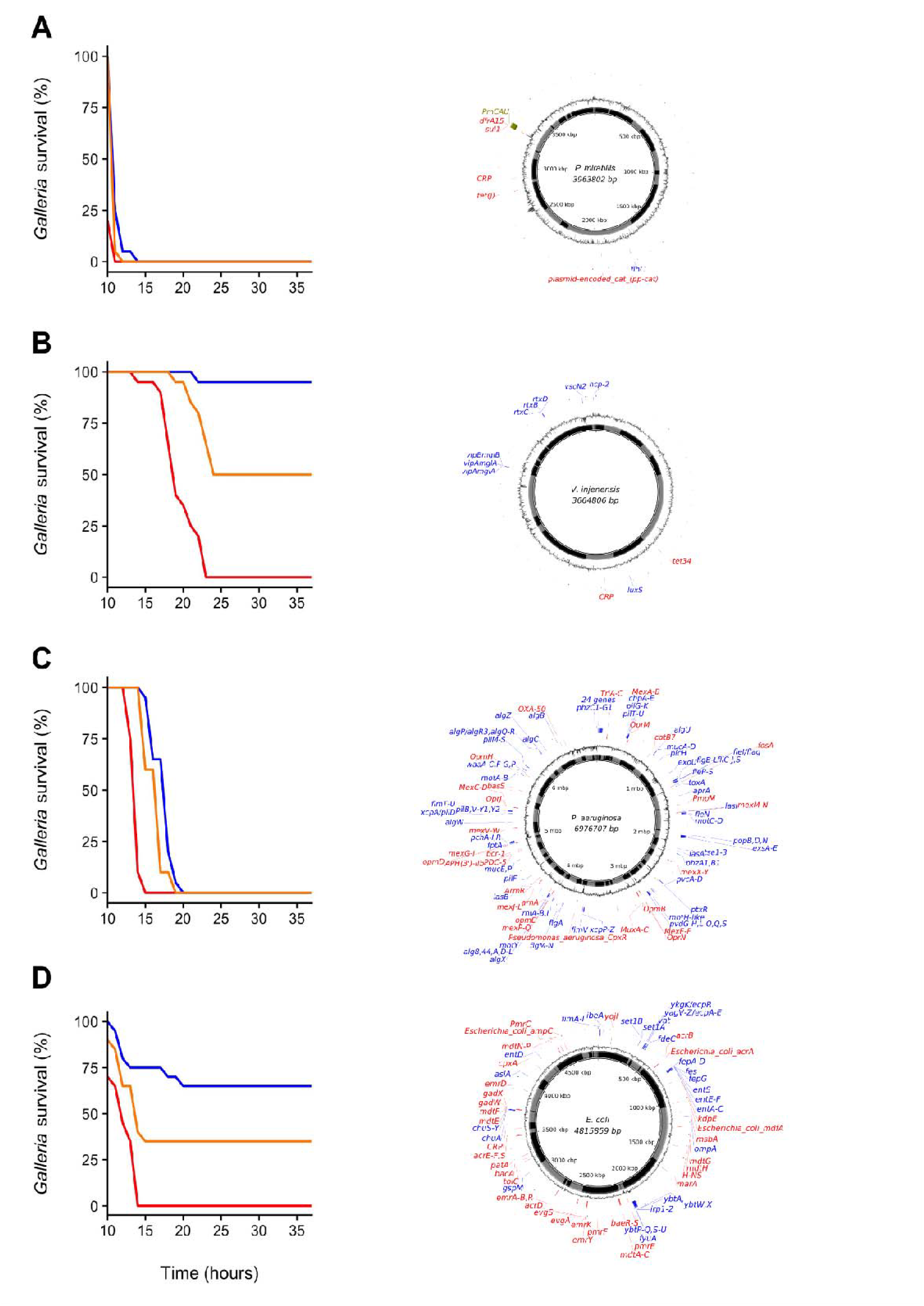
Panels on the left show *Galleria mellonella* mortality after inoculation with individual bacterial clones originally isolated from *G. mellonella* larvae infected with environmental (whole-bacterial community) samples. Groups of 20 *Galleria* larvae were inoculated with 10μL of 1×10^2^ CFU (blue), 1×10^4^ CFU (orange) and 1×10^6^ CFU (red). Panels on the right show clone genome information (species name and genome size (middle), contigs (inner ring; grey and black), GC content (outer ring), virulence genes (blue) and ARGs (red) (≥75% nucleotide similarity used for *P. mirabilis* and *V. injenesis*; ≥90% similarity used for *P. aeruginosa* and *E. coli*; ≥80% coverage criterion for all four species). A: *Proteus mirabilis* (LD_50_= 1×10^2^ CFU) (the genomic island SGI1-PmCAU is indicated in green), B: *Vibrio injenensis* (LD_50_= 1×10^6^ CFU) (note that the absence of a closed draft genome means that contigs are randomly ordered), C: *Pseudomonas aeruginosa* (LD_50_= 1×10^2^ CFU), D: *Escherichia coli* (LD_50_= 1×10^4^ CFU).

The first clone, isolated from estuarine mud (Supplemental Results) was identified as the enteric species *Proteus mirabilis*, most closely related to pathogenic strain HI4320 ^16^ (Fig. 1A). Interestingly, this strain was found to carry a multidrug resistance genomic island (SGI1), first identified in an epidemic *Salmonella enterica* serovar Typhimurium clone in the 1990s ^17^. This island has since been found in *P. mirabilis* isolated from human patients as well as from animals ^18^ but to our knowledge not from *Proteus* strains isolated from natural environments. No virulence genes were found using a 90% similarity cut-off, but several were identified using a 75% cut-off (Table S2). The clone contains several antibiotic resistance genes (ARGs), including the tetracycline efflux protein *TetJ* and *AAC(6’)-Ib7*, a plasmid-encoded aminoglycoside acetyltransferase (90% similarity cut off, Table S3).

The second clone, isolated from beach sand, was found to belong to *Vibrio injenensis*, a recently described species only known from two strains isolated from human patients in Korea ^19^ (Fig. 1B). The UK clone was 99% similar to the type strain M12-1144^T^ and carried 441 genes not present in the Korean strain. Both strains carry the *rtx* toxin operon (Table S4). Only two ARGs, including tetracycline resistance *tet34*, could be identified at a 75% similarity cut off in the UK isolate (Table S5). Unlike the Korean isolate, the UK strain appears to contain a toxin similar to the Zona occludens toxin (Zot) which is known to increase mammalian intestinal permeability ^20^, with highest similarity (74% amino acid similarity) to the fish pathogen *V. anguillarum* ^21^. The isolation of this virulent clone is of particular interest as *Vibrio* species have been identified as high risk emerging infectious pathogens in Europe due to the effects of climate change ^22^.

The third clone *Pseudomonas aeruginosa* (Fig. 1C) isolated from seawater was found to belong to Sequence Type 667, which is represented by four genome-sequenced human pathogens. This clone carries an arsenal of virulence genes (228 at ≥90% nt identity; Table S6) including elastase ^23^ and Type II, III, IV and VI secretion systems. This *Pseudomonas aeruginosa* clone also carries a variety of ARGs (46 at ≥90% nt identity; Table S7), including triclosan- and multidrug efflux pumps and beta-lactamases, including *OXA50* conferring decreased susceptibility to ampicillin, ticarcillin, moxalactam and meropenem, and resistance to piperacillin-tazobactam and cephalotin ^24^.

The fourth clone from estuarine mud was identified as *Escherichia coli* belonging to Phylogroup B2, specifically Sequence Type 3304, represented by three other isolates, from a human patient, a Mountain brushtail possum and one unknown (Fig. 1D). This isolate carries a range of virulence genes (Table S8), including *chuA*, *fyuA* and *vat* known to play a role in uropathogenicity ^25^, *set1A* associated with enteroaggregative *E. coli* ^26^ and ibeA, *OmpA* and *AslA* aiding brain microvascular epithelial cell invasion, known from avian pathogenic- and neonatal meningitis *E. coli* ^27^. This clone contains a range of ARGs, including multidrug- and aminoglycoside efflux pumps, a class C *ampC* beta-lactamase conferring resistance to cephalosporins and *pmrE* implicated in polymyxin resistance (Table S9).

## Discussion

Our study utilized the low-cost and ethically expedient *Galleria* infection model to directly measure the presence of pathogenic bacteria in environmental samples without any prior knowledge of identity. As expected, some samples with low FIB counts contained pathogenic bacteria and some samples with high FIB counts showed low *Galleria* mortality (Fig. S4). We note that of four pathogenic isolates, only one was a coliform and only two were gut-associated bacteria. Two out of the four isolates have not been reported from the U.K. before and potentially represent EIDs. This highlights the fact that transmission risk extends beyond ‘usual suspects’ and includes environmental- and largely uncharacterized strains. Our relatively simple methods can provide a basis for future studies to detect pathogenic bacteria in diverse environments, to ultimately elucidate their ecological drivers and estimate human infection risk.

## Material and Methods

### Sample collection and processing

Eight sampling sites on the Fal estuary and English Channel coast near Falmouth (Cornwall, U.K.) were selected for water and sediment sampling. Four sites were estuarine and four truly coastal (Figure S1). Samples from each site were collected on 21 June 2017 and 6 July 2017. Each sampling effort consisted of collecting water (2 x 50 mL) and upper 1cm layer of sediment (~25 g) from all eight sites within a time span of two hours around low-tide. Samples were collected using sterile 50mL centrifuge tubes. All samples were kept on ice during transport and processed in laboratory within an hour from collection.

Each duplicate water sample was centrifuged at 500 rpm at 4°C for 15 minutes to pellet sediment particles. Supernatants were then transferred to sterile 50mL centrifuge tubes and spun down at 3500 rpm at 4°C for 30 minutes to pellet bacteria. Supernatant were then discarded and the pellet in each tube resuspended in 500μL of sterile M9 buffer. Resuspensions from duplicate samples were then combined in a 1.5 mL Eppendorf tube to obtain a total concentrated water sample of 1mL. Bacteria were extracted from sediments by adding 10g of sediment with 10mL of sterile M9 buffer. Samples were vortexed at 3000 rpm for 2 minutes, and then centrifuged at 500 rpm at 4°C for 15 minutes to pellet sediment particles. The supernatant was then centrifuged as above and resuspended in. 1mL of M9 buffer. All samples were kept at 4°C or on ice at all times.

### Culturable Bacteria

100μL per water or sediment wash sample was plated onto both LB agar (Fisher BioReagents) and Coliform agar (Millipore Sigma) and incubated overnight at 37°C. CFU per plate were counted after 24 hr incubation. Colonies on coliform agar were scored as dark-blue (‘*Escherichia coli*’), pink (‘coliform’), or white/transparent (‘other’). Units are reported in CFU/100mL for water samples and CFU/10g for sediment samples.

### Flow cytometry

Concentrated water and sediment wash samples were analysed for bacterial load using flow cytometry. For each sample (200 μL sample + 20 μL 2.5x SybrGold), 10 μL was analyzed for number of events using low flow rate (14 μL /min) in a BD Accuri C6 Plus flow cytometer (BD Biosciences, San Jose, CA). The BD Accuri C6 Plus is equipped with a blue and red laser, two light scatter detectors, and four fluorescence detectors. Bacteria cells were distinguished from non-biological particles using 533/30 filter in FL1 (indicative of nucleic acid content), 670 LP in FL3 (indicative of chlorophyll content), and FSC (indicative of cell size). Data were analyzed using the BD Accuri C6 software. A non-dye sample was included to control for auto-fluorescence.

### *Galleria mellonella* assay for environmental samples

Final instar waxmoth (*Galleria mellonella*) larvae (~220mg each) were purchased from Livefood UK (http://www.livefood.co.uk), stored in the dark at 4°C, and used within two weeks (all experiments utilized larvae from the same batch). A 100μL Hamilton syringe (Sigma-Aldrich Ltd) with 0.3 x 13mm needle (BD Microlance 3) was used to inject larvae with 10μL of sample into the last left proleg. New sterile needles were used for injection of each sample, and the syringe was cleaned between sample injections with 70% ethanol and sterile M9 buffer. All water and sediment washes (8 locations x 2 timepoints) were assayed in one experiment using 20 *Galleria* per sample. Three negative controls were used for each experiment: a no-injection control (to control for background larvae mortality), a buffer control in which larvae were injected with 10μL of M9 buffer (to control for impact of physical trauma), and a filter-sterilized sample control in which larvae were injected with filter-sterilized samples (0.22 μm syringe filters, Thermo Fisher) to account for any toxic chemicals present in the sample. Before injection, the *Galleria* larvae were separated into Petri dishes in groups of 20 and anesthetized by placing on ice for approximately 30 minutes before injection. After injection, Petri dishes were incubated at 37°C and inspected at 24, 48 and 72 hours post-injection to record morbidity and mortality. *Galleria* larvae were considered infected if they expressed dark pigmentation (melanisation) after inoculation. Larvae were scored as dead if they did not respond to touch stimuli by blunt sterile forceps.

### *Galleria mellonella* assay for individual clones

To isolate and identify responsible pathogens, a subset of samples showing high *Galleria* mortality were inoculated and incubated again as described above. Eight live larvae showing signs of infection (melanisation) were selected at random from each sample group and dissected for haemocoel collection. Before haemocoel collection, the *Galleria* larvae were anesthetized on ice for 30 minutes and the site of dissection was sterilized with 70% ethanol. Sterile micro-scissors were used to remove the last left proleg, and a drop of haemocoel was allowed to exude from the larvae before being collected with a pipette. Approximately 5-15μL of haemocoel was collected per larva and diluted into 500μL of sterile M9 buffer in an Eppendorf tube. Serial dilutions of each sample were plated onto LB- and coliform agar plates and incubated overnight at 37°C. A single colony per sample was picked from these plates and grown for 24 hours in 5mL of LB broth at 37°C at 180 rpm. Overnight cultures were diluted in broth to a turbidity equivalent to a McFarland standard of 0.5 at 625 nm as measured by spectrophotometry (Bibby Scientific Limited, Staffordshire, UK). A serial dilution was plated on LB agar and incubated for 24 hours at 37°Cto obtain cell densities (CFU/mL). Individual clones were used to inoculate groups of 20 *Galleria* larvae with 10μL of 1×10^2^ CFU, 1×10^4^ CFU, and 1×10^6^ CFU. Larvae were incubated as described above and morbidity and mortality was recorded hourly after an initial 10-hour period for a total of 37 hours allowing for construction of survival plots and calculation of the 50% lethal dose value (LD_50_).

### Whole Genome Sequencing

DNA isolation, Illumina HiSeq sequencing and basic bioinformatics was performed through the MicrobeNG program in Birmingham, UK. Vials containing beads inoculated with liquid culture were washed with extraction buffer containing lysostaphin and RNase A, incubated for 25 min at 37°C. Proteinase K and RNaseA were added and incubated for 5 min at 65°C. Genomic DNA was purified using an equal volume of SPRI beads and resuspended in EB buffer. DNA was quantified in triplicates with the Quantit dsDNA HS assay in an Eppendorff AF2200 plate reader. Genomic DNA libraries were prepared using Nextera XT Library Prep Kit (Illumina, San Diego, USA) following the manufacturer’s protocol with the following modifications: two nanograms of DNA instead of one were used as input, and PCR elongation time was increased to 1 min from 30 seconds. DNA quantification and library preparation were carried out on a Hamilton Microlab STAR automated liquid handling system. Pooled libraries were quantified using the Kapa Biosystems Library Quantification Kit for Illumina on a Roche light cycler 96 qPCR machine. Libraries were sequenced on the Illumina HiSeq using a 250bp paired end protocol.

### Bioinformatics

Reads were adapter trimmed using Trimmomatic 0.30 with a sliding window quality cutoff of Q15 (Bolger et al 2014). De novo assembly was performed on samples using SPAdes version 3.7 (Bankevich et al 2012), and contigs were annotated using Prokka 1.11 (Seemann 2014). Kraken was used to indicate the likely species classification of our isolates (Wood and Salzberg 2014). For species with established MLST schemes, the sequence type of each isolate was identified using the Centre for Genomic Epidemiology MLST tool (Larsen et al 2012). Other isolates belonging to the same sequence types were found and accessed using databases Enterobase (Alikhan et al 2018) (*E. coli*) and (Jolley and Maiden 2010) (*P. aeruginosa*). Whole genome alignments were constructed using Progressive Mauve (Darling et al 2010). Percentage similarity was calculated using the number of SNPs found between the query isolate and the reference sequence. In the case of *V. injenesis* where a closed reference genome was not available, nucleotide identity was confirmed using a core genome alignment to closely related draft genome. Core genomes were constructed using Roary 3.12.0 (Page et al 2015).

Differences in gene content (95% nucleotide identity threshold) between the *V. injenesis* reference strain and the *V. injenesis* clone isolated in this study were also identified using Roary 3.12.0. The presence of PCAMU-SGI1 in the P. mirabilis isolate was assessed using BLAST (Altschul et al 1990). Assemblies were screened for antimicrobial resistance related genes and virulence related genes using ABRicate (https://github.com/tseemann/abricat) using 90% and 75% nucleotide identity, and 80% length identity cut offs. The CARD database was used to identify AMR genes (McArthur et al 2013), and the VFDB virulence factor database was used to find putative virulence factors (Chen et al 2015). BRIG version 0.95 was used to represent circular draft assemblies and indicate the position of AMR and virulence genes on each contig (Alikhan et al 2011). Contigs were ordered by whole genome alignment against a reference genome using progressive mauve (Darling et al 2010). The *E. coli* isolate was aligned against K-12 substr. MG1655 (NC_000913.3), the *P. mirabilis* isolate was aligned against HI4320 (NC_010554.1), and the *P. aeruginosa* against PAO1 (NC_002516.2). The *V. injenensis* genome could not be aligned to a closed reference genome and so the contigs could not be ordered or separated by chromosome.

## Data Availability

Trimmed reads and assemblies have been uploaded at the NCBI, Accession: PRJNA473311. Raw data will be deposited into an appropriate database upon acceptance.

## Acknowledgments

We thank Francisca Garcia-Garcia for help with flow cytometry.

## Supplemental Online Material

Contains Supplementary Results, nine Supplementary Tables and four Supplementary Figures.

